# Repeated UV-C exposure alters gibberellin homeostasis and inhibits growth in *Arabidopsis thaliana*

**DOI:** 10.64898/2026.08.22.746412

**Authors:** Adolfo Calvo-Parra Martínez, Theo Lange, Maria J. Pimenta Lange

**Author notes:** **Correspondence:** Theo Lange, and Maria J. Pimenta Lange.

## Abstract

Ultraviolet-C (UV-C) radiation can be highly damaging to plants, yet its effects on gibberellin (GA) homeostasis are not well understood. In this study, we show that short daily UV-C pulse treatments (12 s, 1,200 J m^-2^) applied for seven days reduce plant height and delay flowering in *Arabidopsis thaliana*. Endogenous levels of the GA biosynthesis precursors GA_12_, GA_53_, GA_15_, and GA_24_, the bioactive GA_4_, and the GA catabolites GA_34_ and GA_110_ are all lower in UV-C treated plants than in untreated controls. These changes were accompanied by lower transcript levels of the GA biosynthesis genes *KS, GA13ox1, GA20ox1*, and *GA3ox1*, together with opposing changes in the expression of *GA2ox* genes. Exogenous GA_4_ restores growth in UV-C-treated plants, suggesting that reduced GA availability contributes to UV-C-induced growth inhibition. Consistent with this finding, the GA-signalling mutant *gdella* and the GA-biosynthesis mutants *kao1* and *kao2* show strongly reduced UV-C responses. Together, these findings highlight the importance of GA metabolism and signalling in the developmental response to repeated UV-C exposure, and suggest that exposure regimen influences the dynamics of UV-C-induced hormonal responses.

## Introduction

Ultraviolet (UV) radiation is an important component of the solar spectrum, acting as both an environmental stressor and a developmental signal. It is classified as UV-A (315–400 nm), UV-B (280–315 nm) or UV-C (100–280 nm). While UV-A and UV-B radiation reach terrestrial ecosystems and influence plant growth and development (Ballaré et al., 2011), the amount of UV-C that reaches the Earth’s surface is greatly reduced by atmospheric absorption. UV-C is generally associated with harmful effects in plants, including cellular stress and injury (Danon and Gallois, 1998; Gao et al., 2008). However, its effects are dose-dependent, and low doses can also induce developmental and defence responses. For instance, low-dose UV-C treatments have been shown to promote flowering (Martínez et al., 2004; Takeno, 2016) and enhance plant defences against pathogens like *Botrytis cinerea* (Forges et al., 2018; Vega et al., 2020). Thus, UV-C can act as both a stressor and a developmental and signalling cue. Nevertheless, the mechanisms underlying plant responses to repeated UV-C exposure remain poorly understood.

Gibberellins (Gas) are key regulators of plant growth and development, with well-established roles in various developmental processes (Pimenta Lange and Lange, 2006; Hedden, 2020; Lange and Pimenta Lange, 2020; Hernández-García *et al*., 2021). The coordinated regulation of biosynthesis and signalling controls GA homeostasis. In *Arabidopsis*, GA biosynthesis begins with geranylgeranyl diphosphate (GGPP) in the plastids and proceeds through the activities of *ent*-copalyl diphosphate synthase (CPS) and *ent*-kaurene synthase (KS). This is followed by *ent*-kaurenoic acid oxidases (KAO1 and KAO2), which generate GA_12_ (see Fig. 1). Subsequent reactions catalysed by GA 20-oxidases and GA 3-oxidases produce bioactive C_19_-GAs, including GA_4_. GA activity is further regulated by GA 2-oxidases, which inactivate bioactive and precursor GAs (Hedden, 2020; Lange et al., 2020; Lange and Pimenta Lange, 2020). For instance, C_19_-GA 2-oxidases convert GA_4_ to GA_34_, while some C_20_-GA 2-oxidases act on earlier intermediates, including GA_12_, to produce GA_110_ (see Fig. 1). GA_12_ can also be converted to GA_53_ by GA 13-oxidases. This is a side branch of the pathway that normally contributes relatively little to GA metabolism in *Arabidopsis*. Therefore, changes in the regulation of GA biosynthesis and catabolism can alter GA homeostasis, thereby affecting plant growth and development.

**Figure 1.**
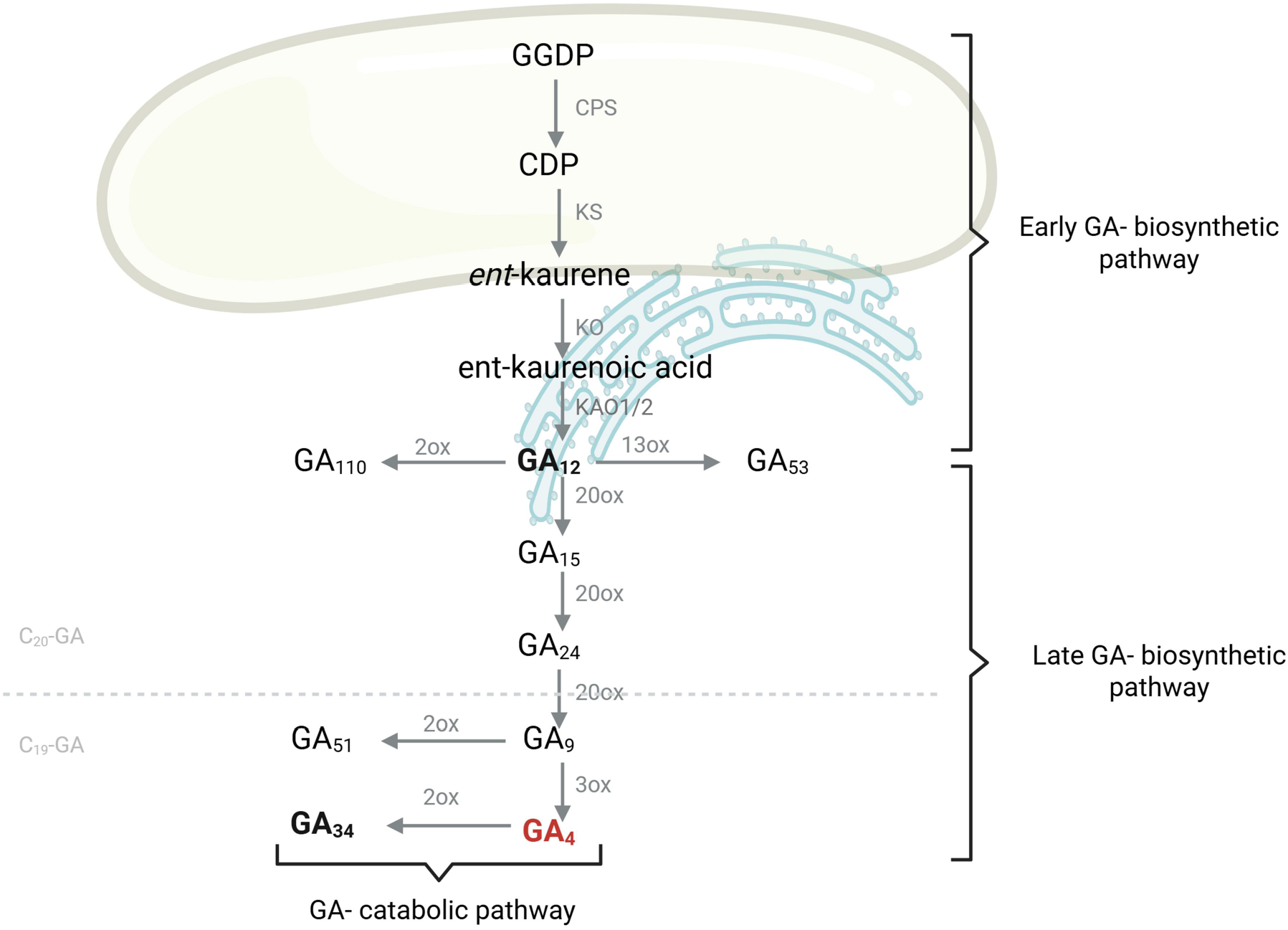
Simplified gibberellin (GA) biosynthetic pathway in Arabidopsis, including the early hydroxylation pathway. Enzymes are indicated in grey: CPS, ent-copalyl pyrophosphate synthase; KS, ent-kaurene synthase; KO, ent-kaurene oxidase; KAO, *ent* kaurenoic acid oxidase; 20ox, GA 20-oxidase; 3ox, GA 3-oxidase; and 2ox, GA 2-oxidase. The central precursor GA_12_, the bioactive GA_4_, and the catabolic product GA_34_ are highlighted in bold. The grey dotted line distinguishes C_20_-GAs from C_19_-GAs.

Recent work by Pimenta Lange *et al*. (2026) showed that a single UV-C pulse induces a biphasic response in *Arabidopsis thaliana*, characterised by an initial reduction followed by an increase in gibberellin (GA) levels, plant height, and flowering time. These findings suggest that the exposure regime is an important determinant of the plant’s response. However, it remains unclear whether repeated UV-C exposure sustains or modifies the initial reduction in GA levels, or how changes in GA metabolism and signalling contribute to the resulting growth phenotype. Given the central role of GAs in regulating growth, GA metabolism and signalling are potential components of the response to repeated UV-C exposure. We therefore investigated how repeated daily UV-C exposure affects plant growth, flowering, GA metabolite levels and the expression of GA metabolic genes. To investigate the functional contribution of GAs, we examined the capacity of exogenous GA_4_ to rescue the phenotype and analysed mutants with defects in GA biosynthesis and signalling.

## Material and Methods

### Plant materials and growth conditions

This study used two Arabidopsis wild-type ecotypes: Columbia (Col-0) and Landsberg erecta (Ler). The global della mutant (*gdella*; *rga-t2, gai-t6, rgl1-1, rgl2-1, rgl3-4*) is in the Ler background (Koini *et al*., 2009). The two mutant lines with impaired *ent*-kaurenoic acid oxidase (*kao1* and *kao2*) are in the Col-0 background. These mutant lines were provided by Dr Patrick Achard (IBMP, Strasbourg, France). The seeds of the wild ecotypes and mutant lines were sown on a soil:vermiculite mixture (2:1, v:v, at 160 g per pot, 92% relative field capacity). The plants were grown in a growth chamber (Aralab FITOCLIMA 600 PLH) at 70% relative humidity and a light-dark cycle of 16 hours light (120 µmol m^-2^s^-1^) and 8 hours dark at 22°C (lights on) and 20°C (lights off).

### UV-C light treatments and phenotype analysis

Unless otherwise stated, two-week-old plants were irradiated with UV-C light (254 nm, 1200 J/m^2^) in a cross-linker (Vilber Lourmat, BIO-LINK/BLX, BLX-254, France) for 12 seconds once per day over a period of one week. The control group (non-irradiated plants) and the UV-C irradiated plants were placed in trays, arranged in a chequerboard pattern and rotated 180° daily within the Aralab chamber to avoid growth effects resulting from the placement. Flowering time was measured as days after sowing, once the inflorescences reached 1.0 cm. Rosette size was measured directly after completion of the treatment in 21-day-old plants using ImageJ software. Plant height was measured of 30-day-old plants, or as specified in the text.

### Exogenous GA_4_ application

Fourteen-day-old Col-0 plants were grown as described above. The plants were either left untreated or subjected to UV-C treatment, as described above. Additionally, half of the untreated plants and half of the UV-C treated plants were sprayed with 10 mL of GA_4_ (10^-4^ M) on days 1, 3, and 5, alongside the UV-C treatment.

### Quantitative analysis of endogenous GAs

Quantitative analysis of endogenous GAs was performed according to the protocol described by Lange et al. (2020). Following the treatments, fresh shoot material of 21-d-old plants was harvested and immediately frozen in liquid nitrogen (N_2_) before being ground. Both the GA quantification and gene expression analyses were performed using this ground material.

### Gene expression analysis

Total RNA extraction, cDNA synthesis and qPCR were performed as described before (Pimenta Lange and Lange, 2015, Pimenta Lange et al., 2026); The primers used for qPCR analysis were described previously (Pimenta Lange and Lange, 2015; Lange *et al*., 2020).

### Statistics

Statistical analysis was performed using Student’s *t*-test with significance levels of *p* < 0.001, *p* < 0.01, and *p* < 0.05 for phenotypic characterization, and of p < 0.05 for GA levels and RT-qPCR data.

## Results

### Repeated short UV-C pulses reduce growth in *Arabidopsis thaliana*

First, we determined the daily UV-C exposure required over a seven-day period to retard growth and development without causing visible damage to the plants. Fourteen-day-old *Arabidopsis thaliana* plants are irradiated with UV-C at 254 nm, receiving 1,200 J/m^2^ for 12, 18 or 24 seconds, once or twice daily (see Supplementary Fig. S1). For subsequent experiments, we selected a 12 s pulse applied once daily because it alters growth without causing severe visible tissue damage. UV-C-treated plants produce shorter inflorescences, flower later, and have smaller rosette leaves than untreated controls (Fig. 2).

**Figure 2.**
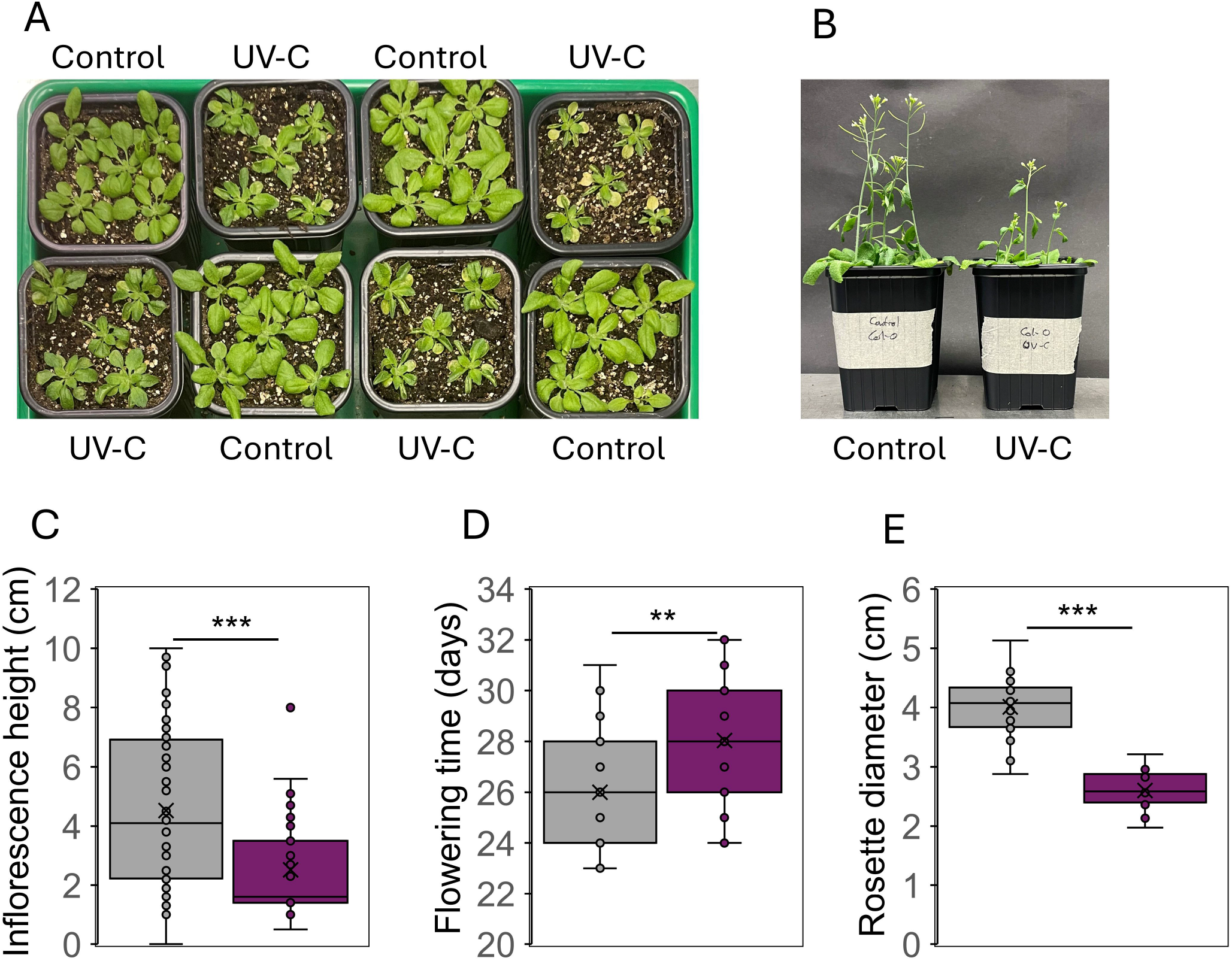
Repeated UV-C exposure inhibits plant development and delays flowering in Arabidopsis. Plants were exposed to a single daily pulse of UV-C for 7 days, as described in the Materials and Methods. **(A)** Phenotypes of control and UV-C-treated 21-day-old plants immediately after completion of the treatment. **(8)** Phenotypes of control and UV-C treated 30-day-old plants. **(C)** Inflorescence height of 31-day-old control and UV-C-treated plants. **(D)** Flowering time of control and UV-C-treated plants. **(E)** Rosette leaf diameter of 21-day-old control and UV-C treated plants. Data are presented as box-and-whisker plots *(n* = 20 plants). Asterisks indicate significant differences between control and UV-C-treated plants *(**p* < 0.01; *\*\*\*p* < 0.001; Student’s t-test).

### Repeated UV-C exposure reduces endogenous GA levels

To determine whether changes in GA metabolism are associated with the UV-C-induced growth phenotype, endogenous GA levels were analysed in plants that were repeatedly treated by UV-C for 7 days. Compared to the control, the UV-C-treated plants show reduced levels of multiple GAs across the biosynthetic pathway (see Figure 3A). This reduction includes the early intermediates GA_12_, GA_53_, GA_15_, and GA_24_, as well as the GA catabolite GA_110_. The bioactive GA_4_ and its catabolite GA_34_ are also present at lower levels in UV-C-treated plants than in the untreated controls.

**Figure 3.**
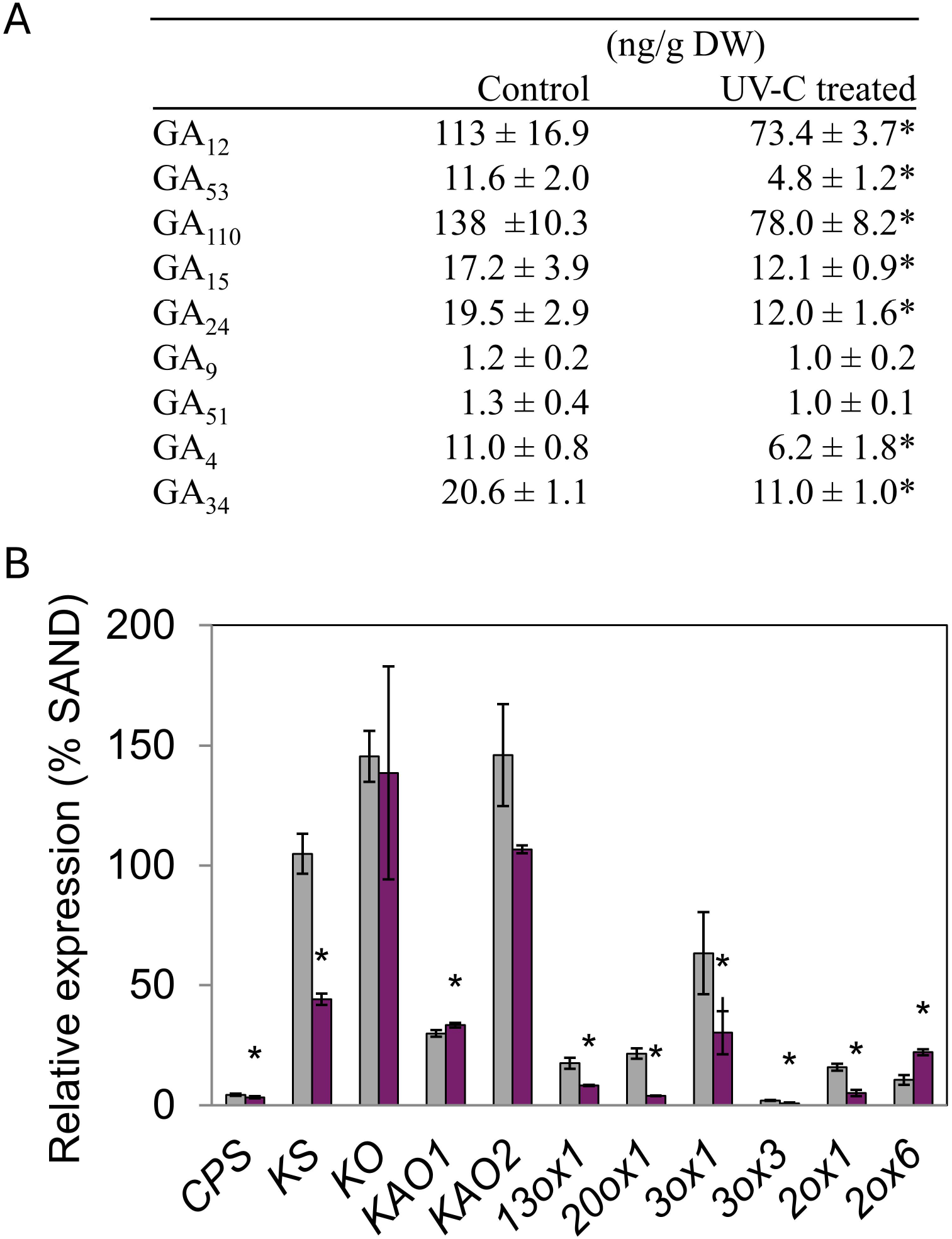
UV-C treatment destabilizes gibberellin homeostasis and GA-related gene expression in Arabidopsis. Plants were either left untreated or exposed to UV-C as described in the Materials and Methods. **(A)** Endogenous GA levels (ng/g dry weight) in 21-day-old plants. Data are presented as means ± SD *(n* = 4). **(B)** Relative expression of GA related genes (% *SAND)*, as described in Materials and Methods. Gray bars represent control plants, and violet bars UV-C-treated plants. Data are presented as means ± Cl *(n* = 3). Asterisks indicate significant differences between control and UV-C-treated plants *(*p* < 0.05; Student’s t-test).

Unlike the biphasic GA response reported following a single UV-C pulse (Pimenta Lange et al., 2026), repeated exposure resulted in sustained lower levels of GA metabolites at the end of the treatment period. Therefore, it appears that the UV-C exposure regime influences GA homeostasis dynamics. To gain a deeper insight into the regulatory process following repeated UV-C treatment of plants, we next analysed the expression of genes related to GA biosynthesis.

### Repeated UV-C exposure alters the expression of genes involved in GA metabolism

To investigate whether transcriptional changes in the GA metabolic pathway could contribute to a reduction in endogenous GA levels, we analysed the transcript levels of genes involved in GA biosynthesis and catabolism in both UV-C-treated and control plants. Following UV-C exposure, the transcript levels of several genes involved in GA biosynthesis, including *KS, GA13ox1, GA20ox1, GA3ox1*, and *GA3ox3*, are half or less than those observed in the control plants (see Fig. 3B). UV-C exposure also altered the expression of genes encoding GA 2-oxidases, albeit not consistently in either direction. Transcript levels of *GA2ox6* are more than twice as high as in the untreated controls, whereas those of *GA2ox1* are less than half (see Fig. 3B). Transcript levels of additional *GA20ox, GA3ox*, and *GA2ox* paralogues remain virtually unchanged (see Supplementary Fig. S2). Overall, these results suggest that repeated UV-C exposure reduces the expression of genes involved in GA biosynthesis, which is consistent with the lower GA levels described above. However, catabolism *via* GA 2-oxidases does not appear to be consistently regulated by UV-C treatment. For instance, transcript levels of *GA2ox6* increase, but GA_34_ levels do not, indicating that transcript levels alone are not sufficient to explain the catabolic profile. Furthermore, the lower levels of catabolites observed in UV-C-treated plants could simply reflect lower levels of available precursors.

### Exogenous GA_4_ rescues UV-C induced growth inhibition

To determine whether reduced GA availability contributes to the UV-C-induced growth phenotype, Col-0 plants were treated with exogenous GA_4_ during repeated UV-C exposure. Previous studies have demonstrated that applying exogenous GA can restore growth in GA-deficient mutants and in plants that have been treated with growth inhibitors (Colebrook et al., 2014).

Application of GA_4_ increases growth in both untreated and UV-C-treated plants, when compared to their respective untreated controls (see Fig. 4A,B). GA_4_ treatment largely restored UV-C-treated plants to the growth of GA_4_-treated controls (see Fig. 4A,B). There is also no significant difference in flowering time between GA_4_-treated UV-C plants and GA_4_-treated control plants (see Fig. 4C). These results support the idea that reduced GA availability contributes to the growth phenotype induced by repeated UV-C exposure.

**Figure 4.**
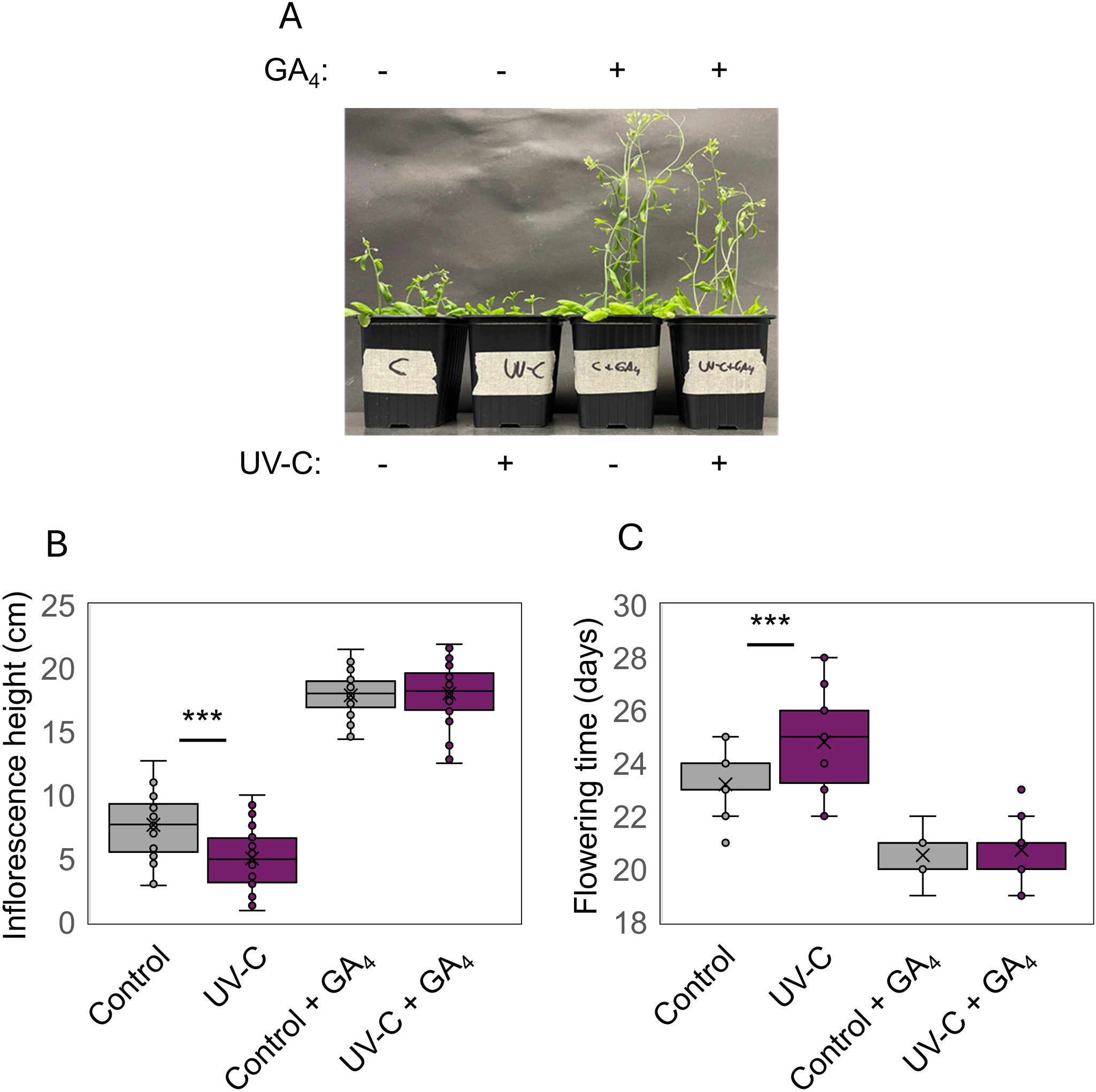
Exogenous GA_4_ rescues developmental changes induced by repeated UV-C exposure in Arabidopsis. Plants were untreated or exposed to UV-C, with or without exogenous GA_4_ application, as described in the Materials and Methods. **(A)** Phenotypes of 30-day-old plants under the indicated treatment conditions. **(8)** Inflorescence height. **(C)** Flowering time. Data are presented as box-and-whisker plots *(n* = 30 plants). Asterisks indicate significant differences between treatments *(***p* < 0.001; Student’s t-test).

### GA-pathway mutants show reduced responses to repeated UV-C exposure

To investigate whether the developmental response to repeated UV-C exposure is regulated by GA biosynthesis and/or signalling, mutant lines affected in the different components of the GA signalling pathway were analysed. These included the GA signalling mutant *gdella*, and the GA biosynthesis mutants *kao1* and *kao2*. Mutant plants and their wild-type controls were subjected to a single daily UV-C pulse for seven days. After this period, they were analysed for inflorescence height and flowering time (Fig. 5A,C,E).

**Figure 5.**
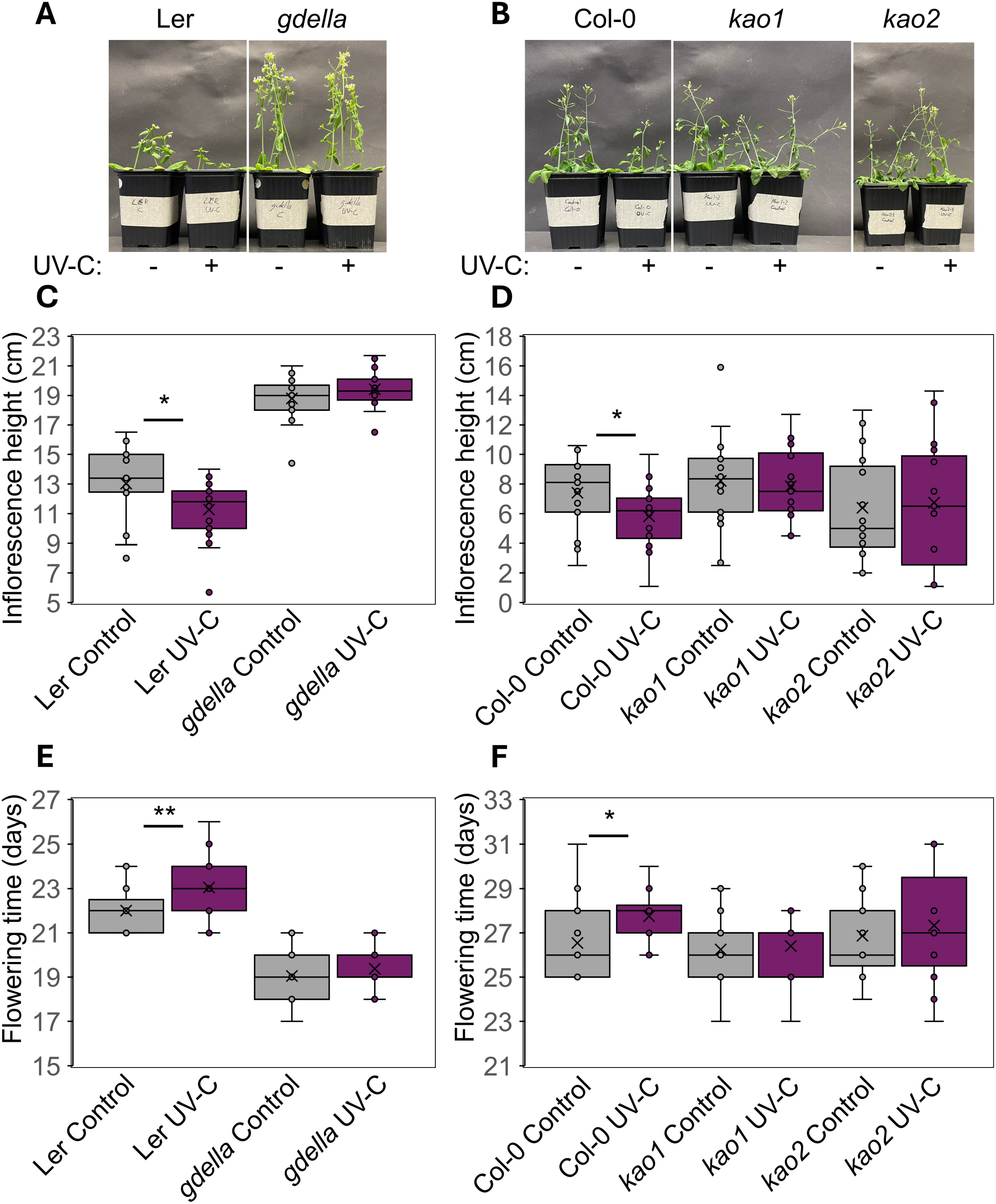
GA signaling regulates flowering time and inflorescence height in Arabidopsis exposed to repeated UV-C treatment. Plants were either untreated(-) or UV-C-treated (+), as described in Materials and Methods. **(A**,**C**,**E)** Phenotypes, inflorescence height, and flowering time, respectively, of *Ler* and *g-della* plants. **(B**,**D**,**F)** Phenotypes, inflorescence height, and flowering time, respectively, of Col-0, *kao1*, and *kao2* plants. **(A**,**B)** Phenotypes of 30-day-old plants. **(C**,**D)** Main inflorescence height of 30-day-old plants. **(E**,**F)** Flowering time. Data are presented as box-and-whisker plots (*n* ≥ 19). Asterisks indicate significant differences between treatments *(*p* < 0.05; *\*\*p* < 0.01; Student’s t-test).

The *g-della* mutant flowered earlier than the Ler wild type under control conditions. However, following repeated UV-C exposure, no significant differences in flowering time or inflorescence height were observed between UV-C-treated and untreated *g-della* plants (Fig. 5).

To further investigate the contribution of GA biosynthesis, *kao1* and *kao2* mutants were analysed. Because KAO1 and KAO2 have overlapping functions in GA biosynthesis and the corresponding single mutants do not exhibit severe developmental phenotypes under standard conditions (Regnault et al., 2014), these mutants provide an opportunity to evaluate the role of early GA biosynthesis to the UV-C response. After seven days of repeated UV-C exposure, no significant difference was observed in either inflorescence height or flowering time between UV-C-treated and untreated *kao1* and *kao2* mutant plants (Fig. 5B,D,F).

Together with the GA_4_ rescue experiments, these results support a role for GA biosynthesis and GA signalling in the developmental response to repeated UV-C exposure.

## Discussion

Our results show that the developmental response to UV-C exposure depends on the regimen used. Whereas a single UV-C pulse has been reported to induce a biphasic GA response (Pimenta Lange et al., 2026), repeated daily exposure resulted in reduced GA metabolite levels accompanied by sustained growth inhibition and delayed flowering. Similar growth-reducing effects of UV-C have been reported in ornamental and crop plants, though the magnitude of these effects and their developmental consequences vary among species and treatment regimens (Bridgen, 2016; Darras *et al*., 2020; Kaliapan *et al*., 2026). The persistence of the retarded growth phenotype, together with reduced levels of multiple GAs, suggests that altered hormonal regulation contributes to the developmental response to repeated UV-C exposure.

The response to repeated UV-C exposure differs from the biphasic response observed by Pimenta Lange *et al*. (2026) after a single one-minute UV-C pulse. In that study, GA levels, plant growth, and flowering time initially decreased but subsequently increased. In contrast, plants exposed to a short UV-C pulse daily for seven days maintain the lower GA levels and exhibit reduced growth and delayed flowering by the end of the treatment period. These differences suggest that the UV-C exposure regimen influences the dynamics of the physiological response. Repeated exposure may prevent or modify the recovery phase observed following a single UV-C pulse. More broadly, variation in UV-C responses across plant species and experimental conditions indicates that the resulting developmental response may depend on the dose, duration, and frequency of exposure (Kaliapan *et al*., 2026).

The metabolite profile also supports the idea of altered GA homeostasis. Following repeated UV-C exposure, the levels of GA_12_ and other GA intermediates (including GA_53_, GA_15_, and GA_24_) were reduced, as were the levels of the bioactive GA_4_ and GA_34_. The reduction in several GA intermediates was accompanied by reduced expression of *KS* and other biosynthetic genes, consistent with reduced biosynthetic capacity. The reduction in GA_34_ is consistent with reduced substrate availability resulting from lower GA_4_ levels, although altered GA2ox activity cannot be excluded because transcript levels do not directly reflect enzyme activity or metabolic flux.

Applying exogenous GA_4_ largely restores both inflorescence height and flowering time in UV-C-treated plants. The ability of exogenous GA_4_ to restore growth indicates that reduced GA activity accounts for a significant part of the developmental phenotype induced by UV-C. These results provide evidence that altered GA homeostasis is functionally linked to the UV-C response. They also imply that GA-dependent mechanisms could be part of a wider response to repeated UV-C stress.

Genetic analyses provide further evidence that multiple components of the GA pathway contribute to the developmental responses. The GA-signalling mutant *gdella* did not show significant changes in flowering time or inflorescence height following UV-C treatment, supporting the idea that GA signalling plays a role in this response. Similarly, the *kao1* and *kao2* mutants did not exhibit a significant response to repeated UV-C exposure. KAO1 and KAO2 catalyse early steps in GA biosynthesis and have overlapping functions in Arabidopsis (Regnault *et al*., 2014). The absence of a UV-C-induced phenotype in these mutants is consistent with the idea that early GA biosynthesis contributes to this response.

Other phytohormones may also contribute to this response. Previous studies have implicated salicylic acid (SA)- and jasmonic acid (JA)-related signalling in plant responses to UV-C radiation (Wang *et al*., 2017; Yao *et al*., 2011). Other hormonal pathways can either regulate GA biosynthesis or interact antagonistically with GA signalling (Unterholzner *et al*., 2015; Xian *et al*., 2024). Pimenta Lange *et al*. (2026) found that *sid2* mutants, which have an impaired ability to synthesise SA, responded similarly to wild-type plants when subjected to UV-C treatment. This suggests that SA signalling alone cannot fully explain the developmental response. Further research is needed to establish whether interactions between GA and other hormonal pathways contribute to the response to repeated UV-C exposure.

Our findings identify exposure regime as an important determinant of the hormonal and developmental response to UV-C. Repeated short UV-C exposure has been shown to alter GA homeostasis in terms of both metabolite concentration and gene expression. This is associated with reduced growth and delayed flowering, as evidenced by the GA_4_ rescue experiment and by the responses of mutants with defects in GA biosynthesis and signalling. These findings suggest that repeated UV-C exposure can reshape hormonal response dynamics and thereby alter developmental outcomes. This approach may provide a sustainable alternative to conventional plant growth regulators in commercial plant production.

## Author contributions

All: investigation and data analysis; writing - original draft; review & editing; MPL, and TL: conceptualization and methodology; supervision.

## Acknowledgements

The authors thank Dr Patrick Achard for kindly providing Arabidopsis mutant seeds. AC-PM would like to dedicate this work to his parents, Maria Luisa Martínez Parra and Fermin Ignacio Villarroya Gil, whose constant support, encouragement and sacrifices made this achievement possible.

## Data availability

The data that support the findings of this study are available from the corresponding authors upon request.

