## Supplementary figures and images for "Repeated UV-C exposure alters gibberellin homeostasis and inhibits growth in *Arabidopsis thaliana*"

### Supplementary Figures

Supplementary Material

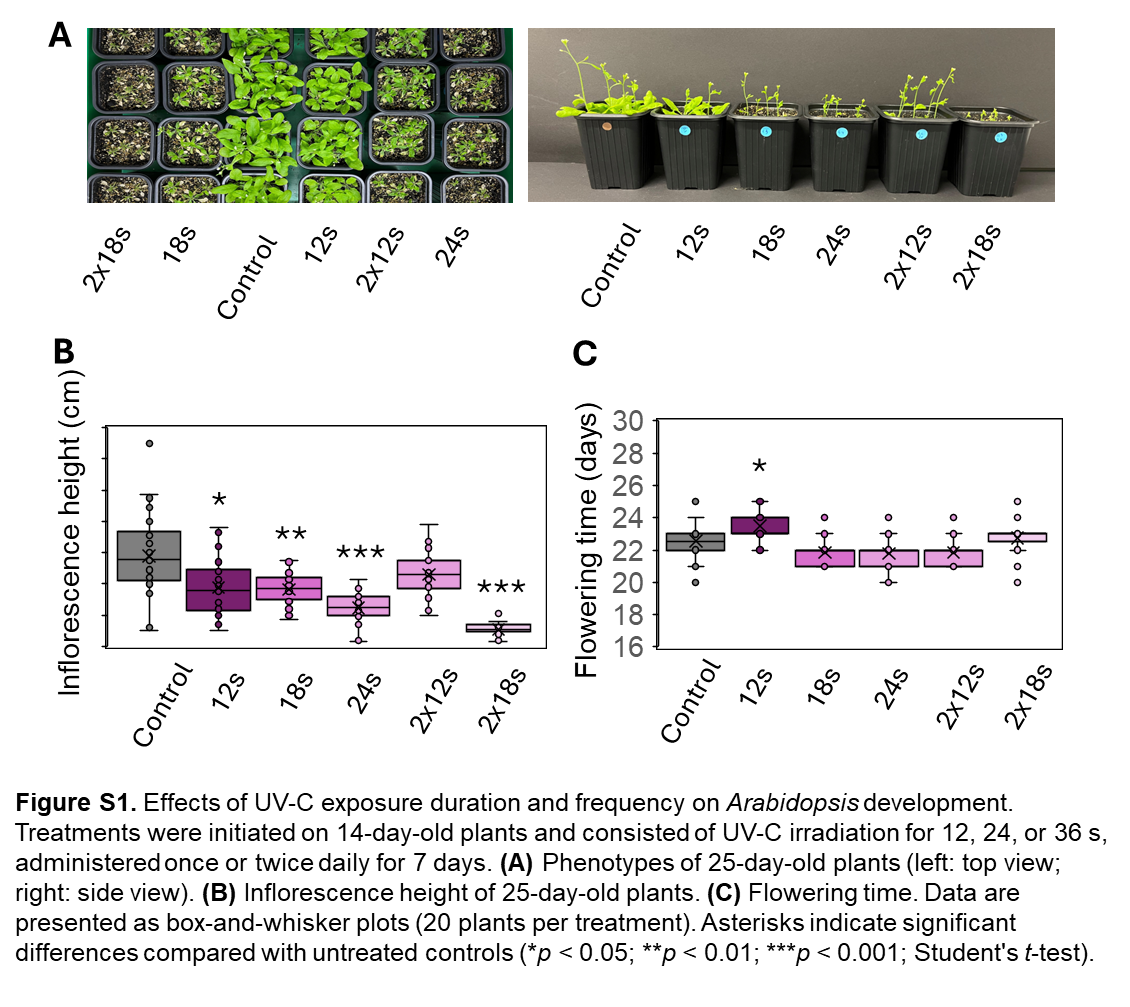

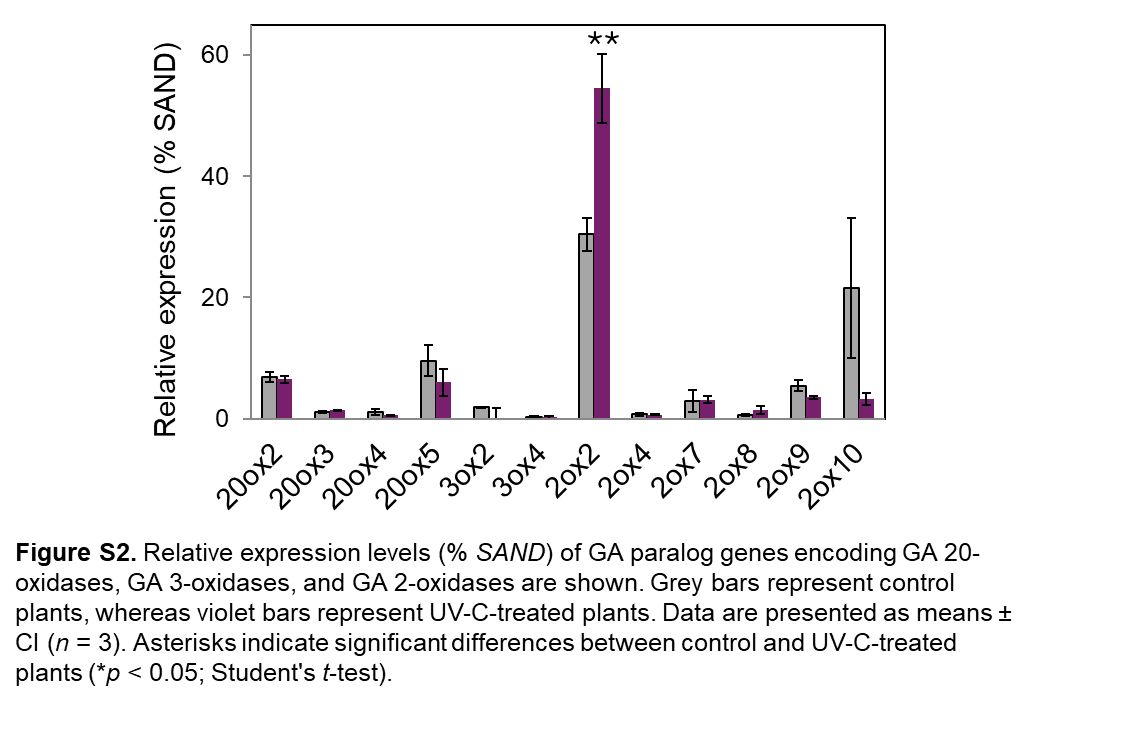
